# Connecting a P300 speller to a large language model

**DOI:** 10.1101/2025.11.06.686984

**Authors:** Mikhail A Lebedev, Anna V Makarova, Daria F Kleeva, Archil I Maysuradze

## Abstract

The advent of large-language models (LLMs) offers a transformative approach for improving the performance of brain-computer interface (BCI) spellers. We propose a novel framework that leverages the contextual understanding of LLMs to compensate for imperfect BCI decoding. Using existing P300 speller data, we simulated a system where users select letters to form words, generating text with characteristic spelling errors. This output is then processed by an LLM, which corrects the errors – a task that becomes more effective when the model considers full-sentence context. Our findings suggest that this synergy can accelerate communication rates by relaxing the need for high single-character accuracy. Beyond speed, integrating an LLM transforms the BCI into an intelligent agent, capable of acting as a discussant and assistant, thereby enriching the user experience.

## Introduction

Brain-Computer Interface (BCI)-based spellers constitute a critical application of BCI technology, developed primarily to restore communication capabilities in individuals with severe motor and speech impairments. Seminal studies with both invasive (Kennedy and Bakay 1998) and non-invasive (Birbaumer et al. 1999) implementations first demonstrated that even completely locked-in patients could regain the ability to communicate by translating cortical activity into computer commands. Among the various paradigms developed, the P300 speller, originally introduced by Farwell and Donchin (1988), has gained widespread adoption due to its relative ease of implementation and reliable, albeit slow, performance. Its original design presents a grid of characters that flash in rapid sequences; the user focuses on a desired character, and the system identifies it by detecting the P300 event-related potential—a positive deflection in the EEG signal occurring approximately 300 ms after a rare, task-relevant stimulus. This foundational paradigm has since undergone extensive modifications to enhance its information transfer rate and accuracy through advanced signal processing, adaptive interfaces, and improved classification algorithms (Fazel-Rezai et al. 2012; Krusienski et al. 2006; Rezeika et al., 2018). Its clinical effectiveness has been robustly validated in populations with amyotrophic lateral sclerosis (ALS) (Sellers and Donchin 2006; Nijboer et al. 2008; Guy et al. 2018) and post-stroke aphasia (Kleih and Botrel 2024). The utility of P300 BCIs extends beyond spelling applications. They have been harnessed for computer gaming (Kaplan et al. 2013; Finke et al. 2009), controlling robotic and prosthetic actuators (Arrichiello et al. 2017; Kaplan et al. 2016), neurorehabilitation and attention training (Arbane et al. 2019), and enabling visuomotor transformations (Bulanov et al. 2021; Syrov et al. 2022).

While reliable, P300-based BCIs suffer from low information transfer rates, typically allowing spelling speeds of only 1-2 words per minute. Research efforts are focused on accelerating operation through optimized stimulus presentation (Lenhardt et al. 2008). Performance can be substantially improved by using invasive signal acquisition methods like electrocorticography (ECoG; Brunner et al. 2011) or stereoencephalography (sEEG; Huang et al. 2020) instead of standard electroencephalography (EEG). Alternatively, a significant speed advantage is offered by SSVEP-based BCIs, which decode responses to rapidly flickering visual stimuli (Chen et al. 2014; Yin et al. 2014; Preetha and Sasikala 2025). This approach has enabled a range of applications beyond spelling, particularly in the control of neural prostheses (Muller-Putz and Pfurtscheller 2007; Ortner et al. 2010).

Recent advancements in LLMs (Minaee et al. 2024; Zhao et al. 2023) are expanding the capabilities of BCI spellers. The integration of language models to aid BCI-based communication is not new (Caria 2025; Mora-Cortes et al. 2014; Speier et al. 2016), with early implementations featuring dictionary-based word completion (Kaufmann et al.; Lee et al. 2011; Ryan et al. 2010). Newer systems like ChatBCI (Hong et al. 2024) and MindChat (Wang et al. 2025) employed more advanced LLMs in P300 spellers to suggest and predict words, thereby accelerating sentence composition. Building on this progress, we propose a framework that leverages the contextual understanding of modern LLMs to compensate for imperfect BCI decoding. Using our previous P300 speller data, we simulated a system where user-selected letters form words containing characteristic spelling errors. An LLM then processes this noisy output to correct mistake – a task that is significantly enhanced when the model utilizes full-sentence context. Our results suggest that this synergistic approach can increase typing rate while offering additional advantages related to communication with an intelligent agent.

## Methods

Here we used the data from our previous study (Kirasirova et al. 2020). The original study was approved by the Samara State Medical University ethics committee (protocol #204, Dec 11, 2019). It involved ten healthy, right-handed 19-year-old males. All participants provided informed consent. EEG signals were recorded using an NVX-36 amplifier with a sampling rate of 250 Hz. Channels P3, Pz, and P4 were sampled using a textile cap with gel-based electrodes.

The experiment tested a P300 speller BCI, where participants focused on target characters in a 4×4 grid (Fig. 1) that flashed randomly. Each character was flashed 10 times per selection. A calibration session was first conducted to train an SVM classifier. Participants then spelled the phrase “JUST DO IT” under two conditions: a standard setup and while wearing an aperture headset. This headset, resembling glasses, restricted the field of view to a single character at a time through a 5mm opening. The order of the two conditions (with and without the aperture) was alternated across four daily sessions, and no significant effect of the condition order was found.

**Figure 1.**
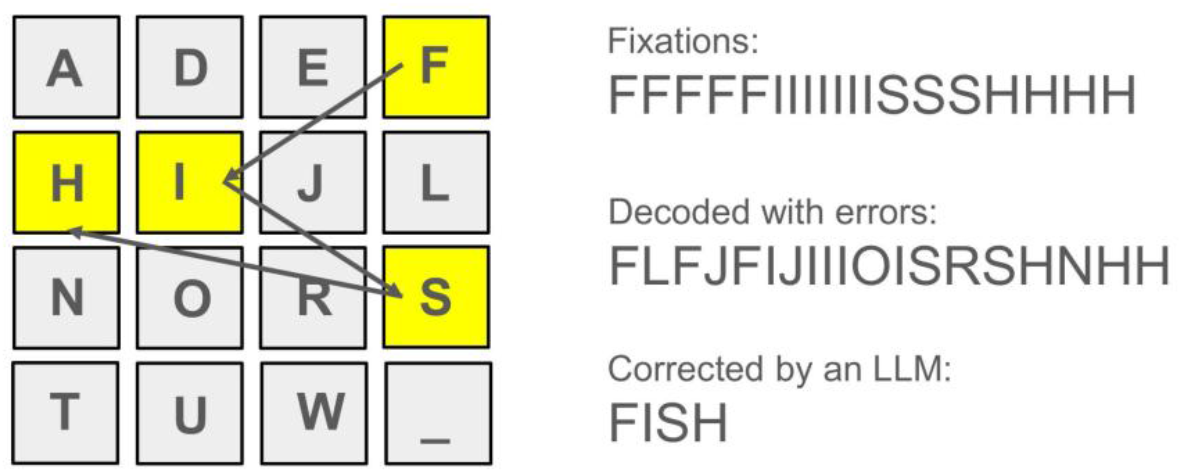
Schematic of the continuous P300 spelling paradigm. A user spells the word “FISH” by gazing consecutively at the corresponding letters on a 4×4 grid. The user aims to fixate on each letter for an optimal number of flashes, but the actual duration can vary naturally (e.g., 5, 7, 3, and 4 flashes, as shown). The BCI decoder translates the EEG signals into a character stream, which may contain errors. This output is then corrected by an LLM to produce the final word.

In this simulation, we modeled participant performance by simulating the spelling of a word (e.g., “FISH”). The simulation assumed a user sequentially fixating on each target letter for a certain number of flashes, guided by an external cue like a periodic background color change. To generate the classifier’s response profile, we pooled epochs from the original recordings as follows: on-target trials were used to represent the attended letter, while random non-target trials were pooled to represent the 15 distractor letters. The actual letter identities from the original data were disregarded.

## Results

Our simulation modeled the following paradigm (Fig. 1): A user views a simplified 4×4 keyboard on a screen while their EEG is recorded. The letters flash according to a predefined schedule (Kirasirova et al. 2020). To select a letter, the user fixates on it for an optimal number of flashes (e.g., five). While the user aims for this target, the actual fixation duration can vary (e.g., 4-6 flashes), and the decoding algorithm accommodates this variability. Since the user can transition between letters fluidly (a process that could be guided by subtle cues like a changing background color) they can focus on spelling entire words rather than individual letters. The system does not require perfect spelling, as an LLM corrects the decoded text. Figure 1 illustrates this process: the user spells “FISH” by fixating F, I, S, and H for 5, 7, 3, and 4 flashes, respectively. The decoder translates the EEG signals into a character stream (which may contain errors), and the LLM then corrects this output to produce the final word: “FISH.”

Figure 2 displays representative EEG responses (Participant 9) to target and nontarget flashes. Panels A and D show the averaged evoked-response waveforms for target (black) and nontarget (magenta) stimuli. To simplify the analysis, we assumed that the participant fixated for the same number of repetitions per character. A comparison of Fig. 2A (5 flashes per character) and Fig. 2D (20 flashes) shows a clear increase in the signal-to-noise ratio with longer fixation durations. We quantified the response similarity to the target response using the correlation coefficient, r. This metric measures the correlation between the average EEG trace for a given character and the target template, which was derived from the average of 100 target stimulus responses. The frequency distributions of r are shown for 5 flashes (Figs. 2B, C) and 20 flashes (Figs. 2E, F). The distributions for nontarget and target responses are shown in magenta and black, respectively. As the number of repetitions per character increases from 5 to 20, the separation between the target and nontarget distributions becomes more distinct, indicating improved discriminability.

**Figure 2.**
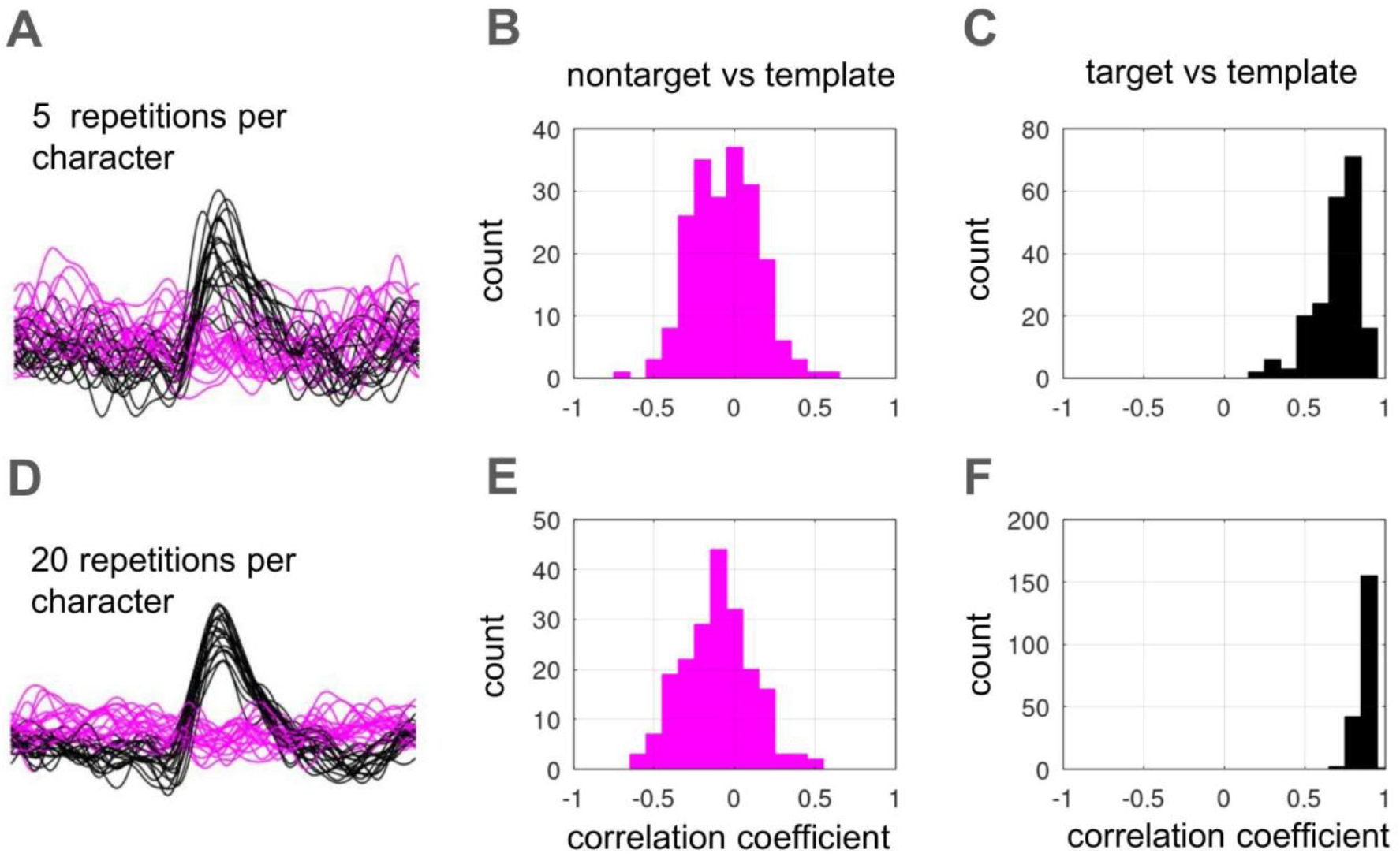
Improved target discriminability with increased stimulus repetitions. A, D: Averaged EEG waveforms from a representative participant (Participant 9) in response to target (black) and nontarget (magenta) flashes for 5 and 20 repetitions, respectively. The signal-to-noise ratio increases with more repetitions. B, C, E, F: Frequency distributions of the correlation coefficient, which quantifies the similarity of individual responses to a target template. Target (black) and nontarget (magenta) distributions show greater separation with 20 flashes, indicating improved classification.

While various classification algorithms could be used, we used a simplified method that converted evoked potentials into letters using r as a metric. The value of r was computed for each of the 16 possible characters; the character upon which the participant was fixating yielded the highest r-value, and the decoder selected this maximum. For simplicity, we assumed that the decoder had already identified the time periods of distinct gaze fixations. In Figs 3 and 4, the evoked potentials are presented in a color-coded matrix, with rows showing average EEG traces for different characters. The user’s intended characters are distinctly evident with 20 repetitions (Fig. 3), whereas the pattern is less discernible with only 5 repetitions (Fig. 4), though the word (“FISH”) is accurately identified in both cases.

**Figure 3.**
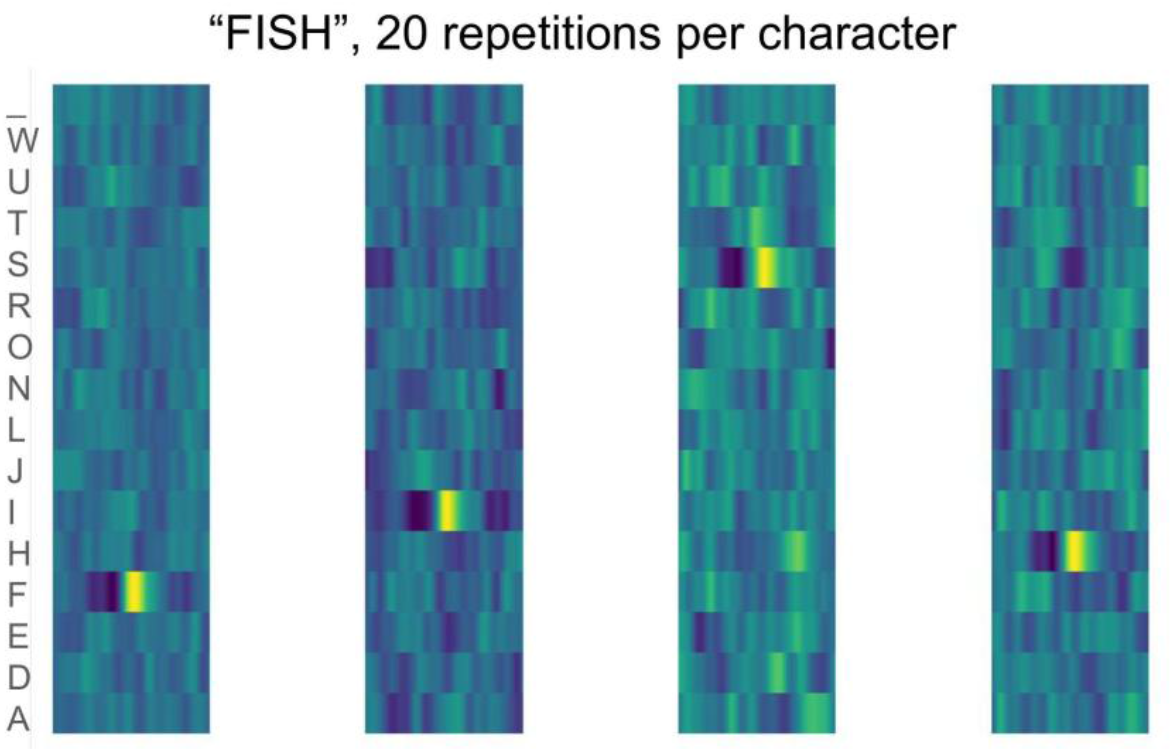
Decoding of the word “FISH” from evoked potentials. Color-coded matrices show the character-specific evoked potentials. A correlation-based classifier (selecting the character with the highest r-value) successfully identified the word using 20 repetitions per character.

**Figure 4.**
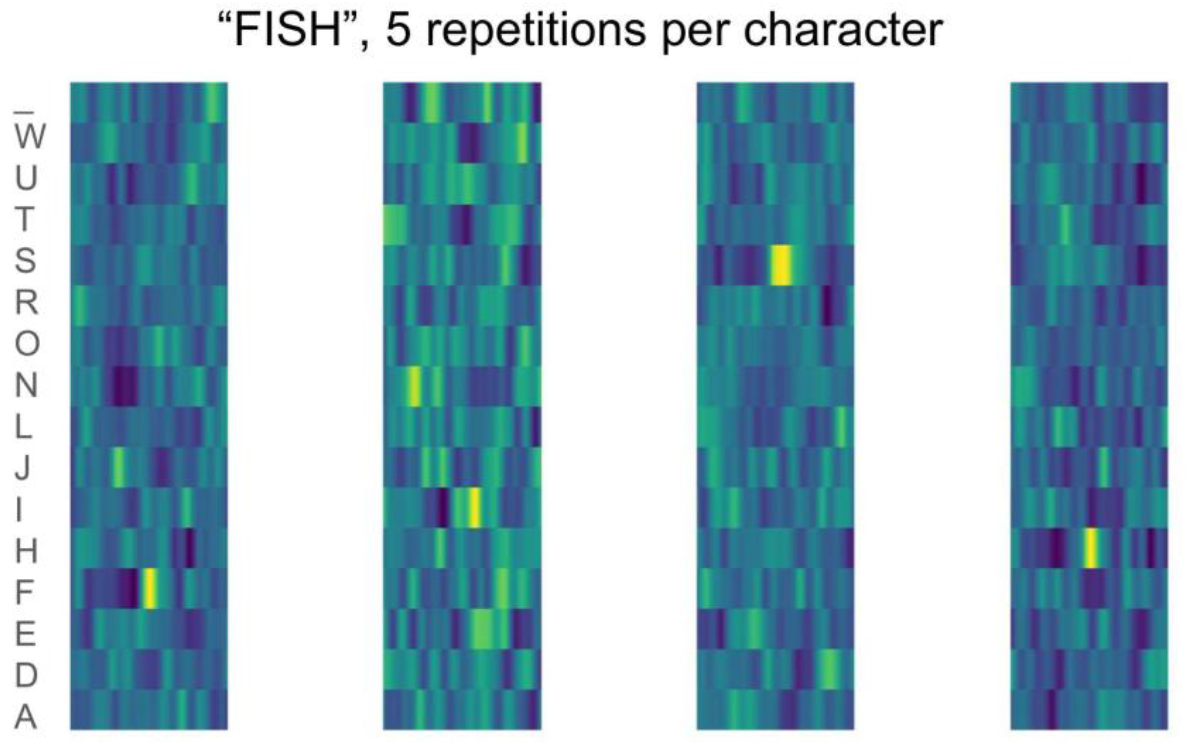
Decoding of the word “FISH” using 5 repetitions per character. Conventions as in Fig. 3.

Figure 5 displays the r-values for decoding the word “FISH” using 2, 5, and 10 repetitions per character (Panels A, B, and C). In these color-coded matrices, columns represent the consecutive letters of the word and rows represent keyboard characters. With 10 repetitions (Fig. 5C), the spelled letters are clearly discernible, while fewer repetitions produce noisier patterns. The results from 10 decoding attempts, shown below the colorplots, demonstrate that the word was correctly decoded in 40%, 70%, and 100% of trials for 2, 5, and 10 repetitions, respectively.

**Figure 5.**
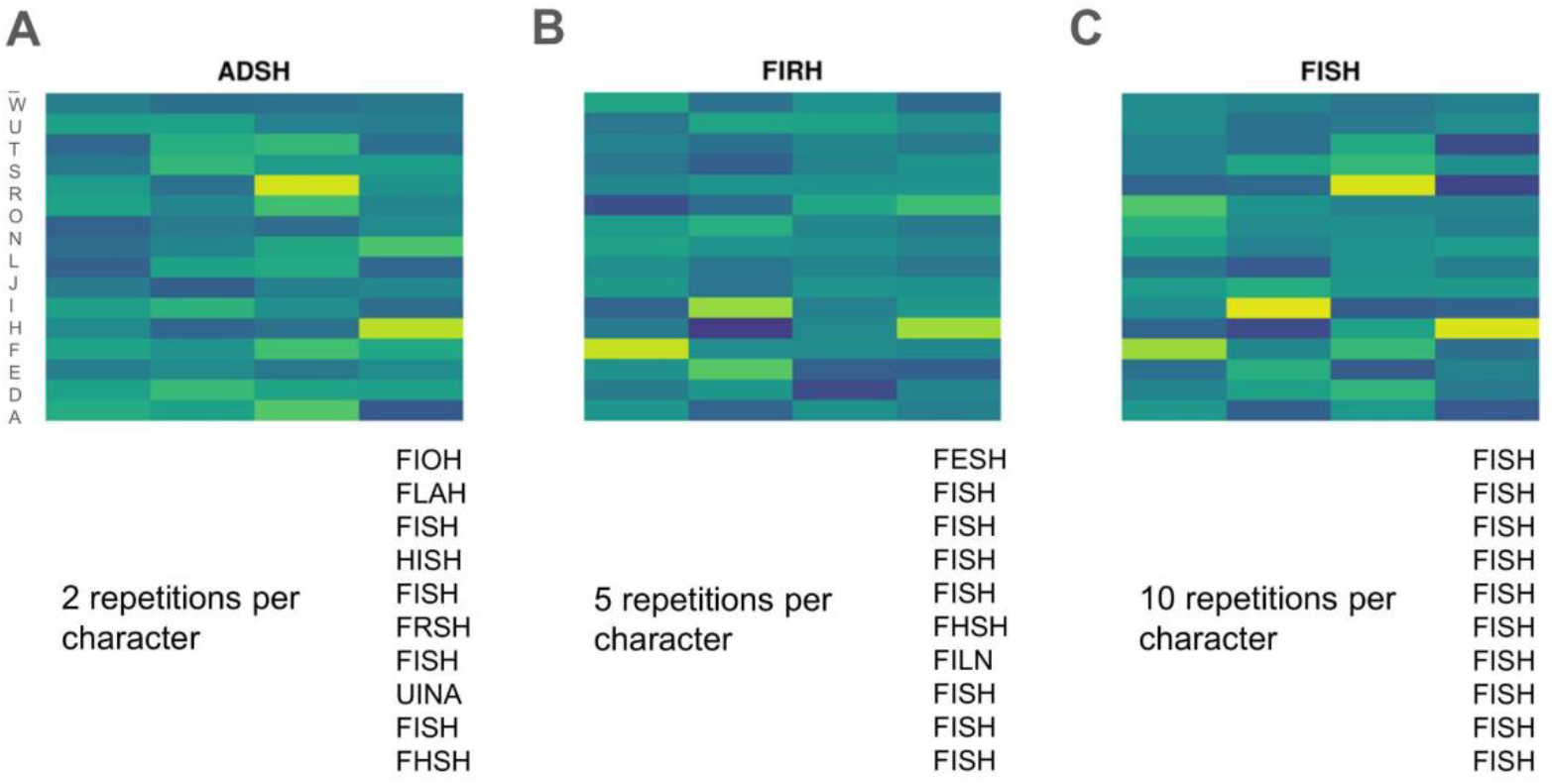
Decoding of the word “FISH” with different character repetition counts. R-value matrices for spelling “FISH” with 2 (A), 5 (B), and 10 (C) repetitions per character. Rows: keyboard characters; Columns: target letters. The correct spelling is visually clear with 10 repetitions. Decoding accuracy (bottom) increased from 40% to 100% across conditions.

This same method of visualization is extended to a more complex example in Fig. 6, which illustrates the decoding of the sentence “HE WASHED HIS HANDS WITH THE FRESH WATER” using 5 repetitions per character. The initial, unprocessed output from the EEG speller was the character sequence “HT WASHED HIS HAN S WITILTHE THE FRESH WATER”. Following this, the resulting text string was passed as an input to three separate LLMs— specifically ChatGPT, DeepSeek, and Grok—for post-processing. All three LLMs were independently and successfully able to reconstruct the original, intended sentence from the noisy output.

**Figure 6.**
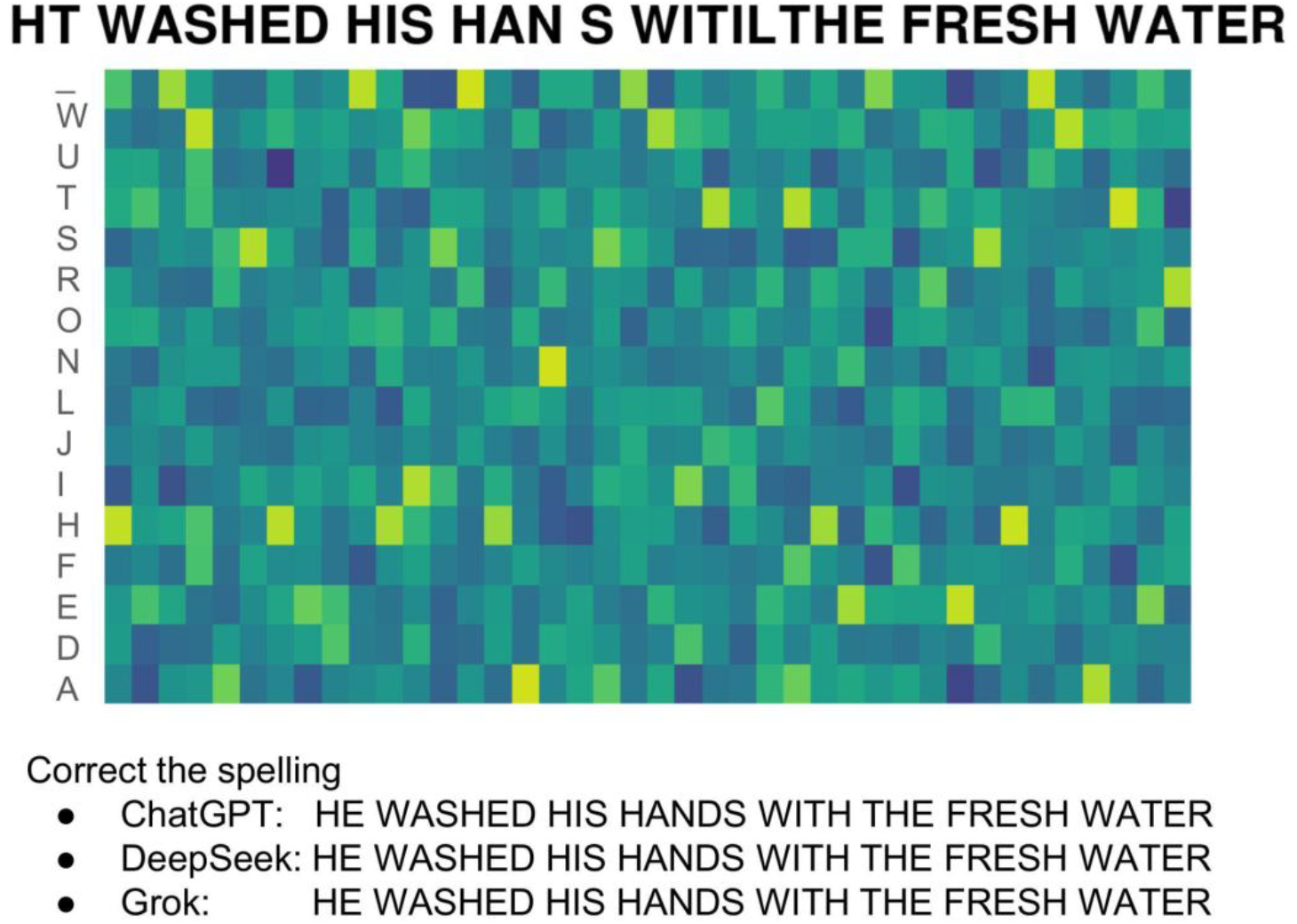
Sentence decoding with LLM post-processing. The r-value matrix for decoding the sentence “HE WASHED HIS HANDS WITH THE FRESH WATER” using 5 repetitions per character. The initial EEG speller output was the erroneous sequence “HT WASHED HIS HAN S WITILTHE THE FRESH WATER”. When this output was processed by three large language models (ChatGPT, DeepSeek, and Grok), all three successfully reconstructed the original, intended sentence.

## Discussion

The present study successfully provides a proof-of-concept demonstration that integrating an LLM with a P300 speller can effectively compensate for imperfect BCI decoding, thereby enhancing communication speed and reliability. Simulation results, based on empirical EEG data, confirmed that while lower numbers of character repetitions led to noisier output and more spelling errors in the raw BCI stream, subsequent processing by an LLM corrected these mistakes.

We did not introduce dramatic modifications to the basic P300 paradigm (Farwell and Donchin 1988; Fazel-Rezai et al. 2012; Krusienski et al. 2006; Rezeika et al., 2018) but rather proposed several practical improvements. First, we shifted the core philosophy from requiring perfectly spelled individual letters to spelling entire words as discrete entities. In this approach, the user views a typical P300-speller display but proceeds through a word at a comfortable, self-paced rhythm. They focus on individual keys for a variable number of flashes before moving to the next. The decoder’s role is to identify the boundaries of these fixation epochs. Users could receive feedback of their performance, for instance, through keys highlighted with a luminance proportional to their decoded probability. This would reinforce that longer fixations yield better decoding. However, to prevent slow typing speeds, our design allows users to find a balance between speed and accuracy. It tolerates spelling errors at the word level, which are subsequently corrected by a post-processing Large Language Model (LLM) that ensures accuracy at the sentence and contextual level once enough text is available.

While we initially tested these concepts with P300 data (Kirasirova et al., 2020), they are equally applicable to an SSVEP approach (Chen et al., 2014; Yin et al., 2014; Preetha & Sasikala, 2025), which may offer superior speed. Furthermore, while EEG recordings inevitably contain oculomotor components often treated as artifacts (Schlögl et al., 2007), these are legitimate electrophysiological signals. A practical decoder could leverage them to identify periods of stable fixation and even aid in predicting the next keyboard fixation point.

The present work utilized a basic template-matching approach for P300 decoding. Modern practical implementations would benefit from more advanced classifiers, a diverse set of which have been established in the field (Krusienski et al. 2006; Lotte et al. 2018). A notable recent advancement is the application of transformer architecture, which show considerable promise for enhancing P300 classification accuracy (Gomes de Novais et al. 2024; Hong and Najafizadeh 2024; Li et al. 2022).

As to the integration of a P300 BCI with an LLM, it is clear from our assessment that LLMs are perfectly suited to work with the text generated by a speedy but imprecise BCI and convert this text with errors into clearly written sentences. Several ways to integrate a P300 speller with an LLM has been recently proposed (Babu et al. 2025; Caria 2025; Hong et al. 2024; Wang et al. 2025). The future of BCI-LLM systems will not be limited to improved text decoding, as they are poised to offer a whole range of interactions that disabled people need. One direction for the future is the creation of LLM-incorporating BCIs that act as a cognitive co-pilot that leverages the LLM’s generative and contextual capacities to decode the user’s intent, reducing the long typing to a few high-level conceptual commands. Such hybrid systems could enable control over the user’s environment, including smart home devices and browsing the internet. Recognition of emotions could be also helpful to improve understanding of context and communications (Torres et al. 2020). And of course, the EEG-based systems could be also an intermediate step toward high capacity invasive BCI (Musk 2019) which will also need integration with LLMs to improve efficiency.

## Acknowledgements

This work was supported by the The Ministry of Economic Development of the Russian Federation in accordance with the subsidy agreement (agreement identifier 000000C313925P4H0002; grant No 139-15-2025-012).

